# Angular speed should be avoided when estimating the speed-curvature power law in movement

**DOI:** 10.1101/2022.06.27.497695

**Authors:** Adam Matić

**Affiliations:** Instituto de Neurociencias CSIC-UMH, Alicante, Spain

**Keywords:** motor control, speed-curvature power law, geometry, kinematics

## Abstract

The speed-curvature power law is one of the most studied constraints in biological movement. In many types of movements, there is a strong relationship between instantaneous speed and local curvature. For example, in elliptical trajectories, tangential speed is proportional to curvature raised to the power -1/3, (V∼C^-1/3^). This phenomenon is known as the “one-thirds power law” and is generally considered to be mathematically equivalent to the “two-thirds power law” that describes the relationship between angular speed and curvature (A∼C^2/3^); the two formulations are used interchangeably. However, in this paper, analysis of empirical and synthetic data demonstrates that using angular speed instead of tangential speed to estimate the power law tends to result in much stronger correlations, impacting the interpretation of the strength of the relationship, and therefore the existence of the law. Further analysis shows that angular speed and curvature are often trivially correlated, since angular speed is not a purely kinematic variable and depends on curvature. In conclusion, two forms of the law are not equivalent, angular speed should be avoided when expressing the speed-curvature power law.

## 1. INTRODUCTION

The relationship between instantaneous speed and local curvature in human movement is a classic, well-studied phenomenon in the field of motor control. Its historical roots can be dated to the observations of Binet and Courtier (1893) and Jack (1895) that in *natural handwriting* - natural being not too slow nor too fast - the speed of the pen tends to be lower in curved segments of the path and higher in the straight segments. Lacquaniti and colleagues (1983) proposed that the relationship can be expressed as a power law, and reported its validity for a large class of hand movements. The law has since been observed also in human locomotion, speech, eye movements, (for a review, see Zago, Matič et al. 2017). It has also been observed in the trajectories of animals (Zago et al, 2016, James et al. 2019, Dagenais, et al 2021), and has been used a criterion for considering simulations of movement sufficiently life-like (Loveless et al 2019, Matič et al. 2021). Taking inspiration from animal movement, roboticists incorporate the law for trajectory planning for robots (Shafiei et al. 2015; Rybarczyk and Carvalho 2019).

The speed-curvature power law is expressed in several ways, involving either tangential speed or angular speed as kinematic variables and curvature or radius of curvature as geometric variables. Researchers often explicitly state that the different formulations of the power law are mathematically *equivalent* (e.g. Lacquaniti et al. 1983; Viviani and Cenzato, 1985; Wann et al. 1988; Viviani and Schneider, 1991; Schaal and Sternad, 2001; Vieilledent et al. 2001; Ivanenko et al. 2002; Perrier and Fuchs, 2008; Maoz et al. 2009; Tesio et al. 2011; Huh and Sejnowski, 2015; Karklinsky et al. 2016; Zago, Matić et al. 2017; Rybarczyk and Carvalho 2019; Matić and Gomez-Marin 2020).

Indeed, examining the algebraic relationships shows that the exponents in different formulations of the power law are mathematically related. Namely, with angular speed A, curvature C, and a constant k (related to average speed, also called the velocity gain factor), the “AC power law” is:

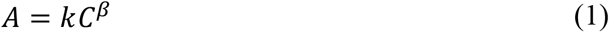

With tangential speed V, since A=VC, the “VC power law” is:

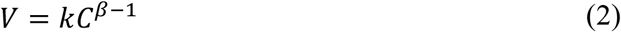

If instead of curvature C, we use the radius of curvature R = C^-1^, the exponent of the power law is: *A=kR*^*−β*^or *V=kR*^1*−β*^. See also Figure 1 for definitions of variables.

**Figure 1.**
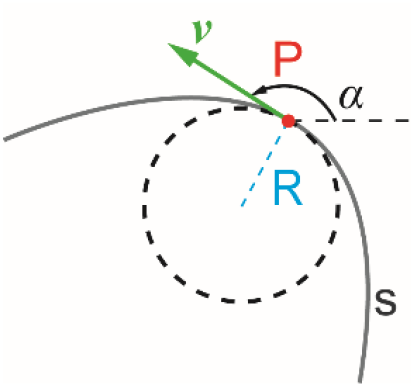
For a particle P moving along the path s, speed (V) is defined as the magnitude of the (tangential) velocity vector,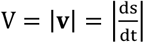, while angular speed (A) is the absolute rate of change of the direction of velocity,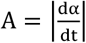, and curvature (C) is the reciprocal of the radius of the osculating circle, or the rate of change of direction with respect to arc-length,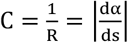. From this we see that, naturally, A=VC.

Apparently, following equations (1) and (2), and delaying the analysis of the correlations, the AC and VC power law are equivalent, since the exponent of the VC power law can always be calculated from the exponent of the AC power law. Indeed, in drawing ellipses and scribbling movements, the exponent of the AC power law (equation 1) is empirically found to be β≈2/3 (Lacquaniti et al. 1983), a phenomenon known as the “2/3 power law”. If we choose to analyze the same trajectories with the VC power law (equation 2), the exponent is near -1/3, and the phenomenon is sometimes called the “1/3 power law”. Regardless of the name used, most published research uses tangential speed and curvature, the VC power law, as expressed in equation (2). However, the repeated statements to the equality of the laws and the explicit use of angular speed are frequent enough to warrant scrutiny (e.g. Lacquaniti et al. 1983, Ivanenko et al 2002; Zago, Matić et al. 2017; James et al. 2019).

Crucially, the mathematical relationships in (1) and (2) show an equivalence of exponents between relationships, but not the equivalence of the *strength* of the underlying relationship.

In the research on the power law of movement, power functions (1) and (2) are typically converted to linear functions by taking a logarithm of both sides. Next, the exponents β, the speed gain factor k and the strength of the relationship, expressed as the coefficient of determination r^2^ are estimated by linear regression from models:

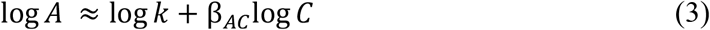

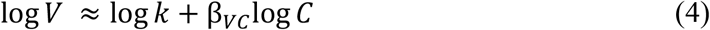

In linear regression, the coefficient of determination (r^2^), shows how much variance in the outcome is explained by the predictors, expressing the strength of the linear relationship. When there are only two variables in a regression, the r^2^ is the square of the correlation coefficient r (Pearson’s or other). More generally, it is calculated as r^2^ = 1 – RSS/TSS where RSS is the sum of squares of residuals and TSS the total sum of squares of the outcome. If the relationship between variables has a high r^2^ (near 1), one can predict with high accuracy the value of the outcome variable from the values of the predictors.

## 2. EMPIRICAL DATA: Tracking elliptic targets at different frequencies Methods

The following is an analysis of pen trajectories recorded in a target tracking experiment (Matič, 2022). Participants (N=3) were seated, looking at a computer monitor, while holding an electronic pen in their dominant hand on a graphics tablet (Wacom Intuous). There was a small circular target displayed on the screen. The pen determined the position of a circular cursor on the screen, and the task of the participant was to track the target with the cursor. Targets always moved in an elliptical pattern. Target trajectories varied across trials (i) in 9 different frequencies, from slow rhythms to fast, and (ii) in 3 different speed profiles: constant speed (β_VC_=0), ‘natural speed’ (β_VC_=-1/3) and ‘extra slowing’ (β_VC_=-2/3). Each participant (three males, age 30-36) performed 27 trials of target tracking, each trial lasting 16 seconds, presented in a random order.

The recorded trajectories were smoothed by a second-order, low-pass Butterworth filter with a cutoff at 10Hz, and differentiated to estimate speed and curvature. The power law exponents β and the coefficients of determination r^2^ were estimated using linear regression. Conformity to the power law was chosen at an arbitrary point of r^2^ ≥ 0.75, considering that pure gaussian noise can produce a fit of up to r^2^ ≈0.6 (Maoz et al, 2005).

All of the data and the Python code for analysis and generating the figures are available in an online repository: https://github.com/adam-matic/AngularSpeedPowerLaw.

## Results

The plot of the VC power law across frequencies (Figure 2) and the plot of the AC power law across frequencies (Figure 3) show that the values of exponents are equivalent; for each trajectory the following equality holds: β_AC_= β_VC_ + 1. However, there is a dramatic difference in the coefficients of determination (r^2^). For the VC power law, only the trajectories performed at fast rhythms of f ≥ 0.94 conformed to the power law with the coefficient of determination of r^2^ ≥ 0.75. On the other hand, angular speed is strongly correlated to curvature across the entire frequency range, with only a few exceptions of the power law estimates with r^2^ < 0.75, and even then, the lowest value is a relatively high r^2^ ≈ 0.5.

**Figure 2.**
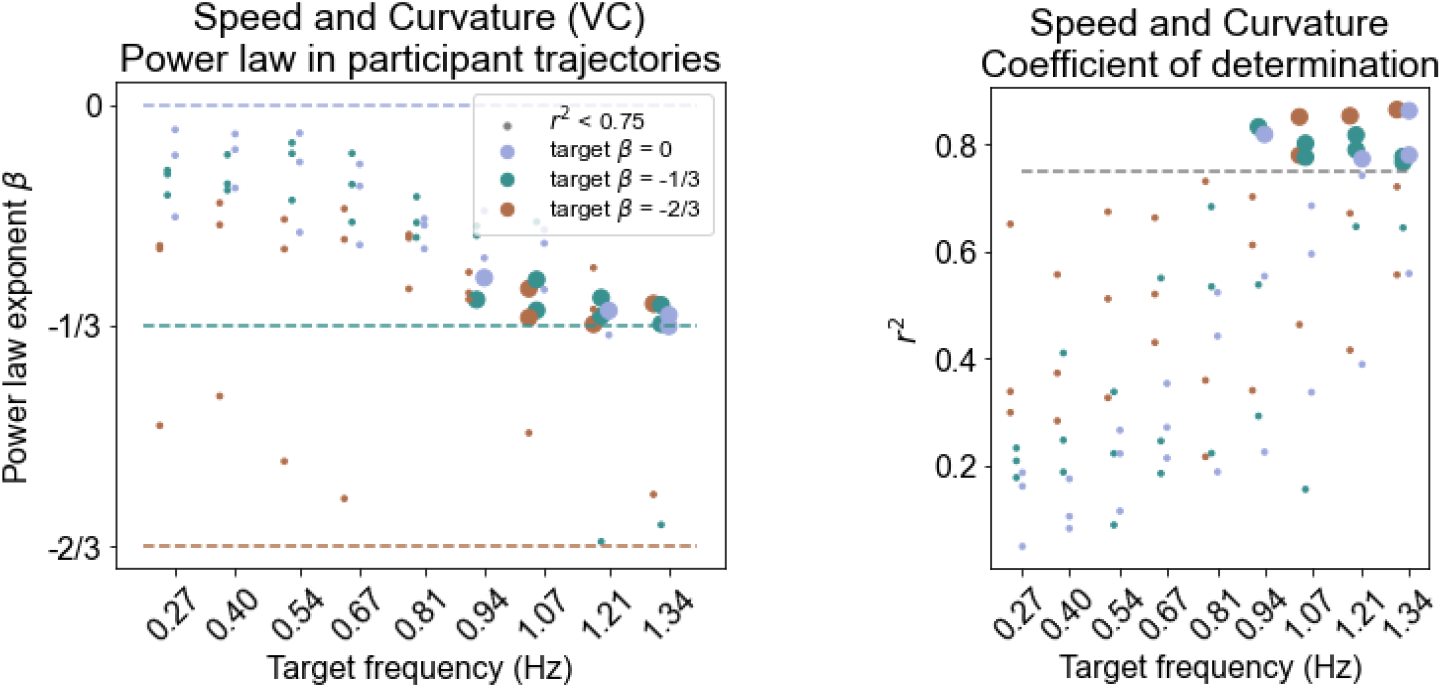
Using tangential speed and curvature to estimate the speed-curvature power law, only high-frequency participant trajectories (freq ≥ 0.94) conform to the power law with a coefficient of determination r^2^ ≥ 0.75. Many trajectories have a low r^2^.

**Figure 3.**
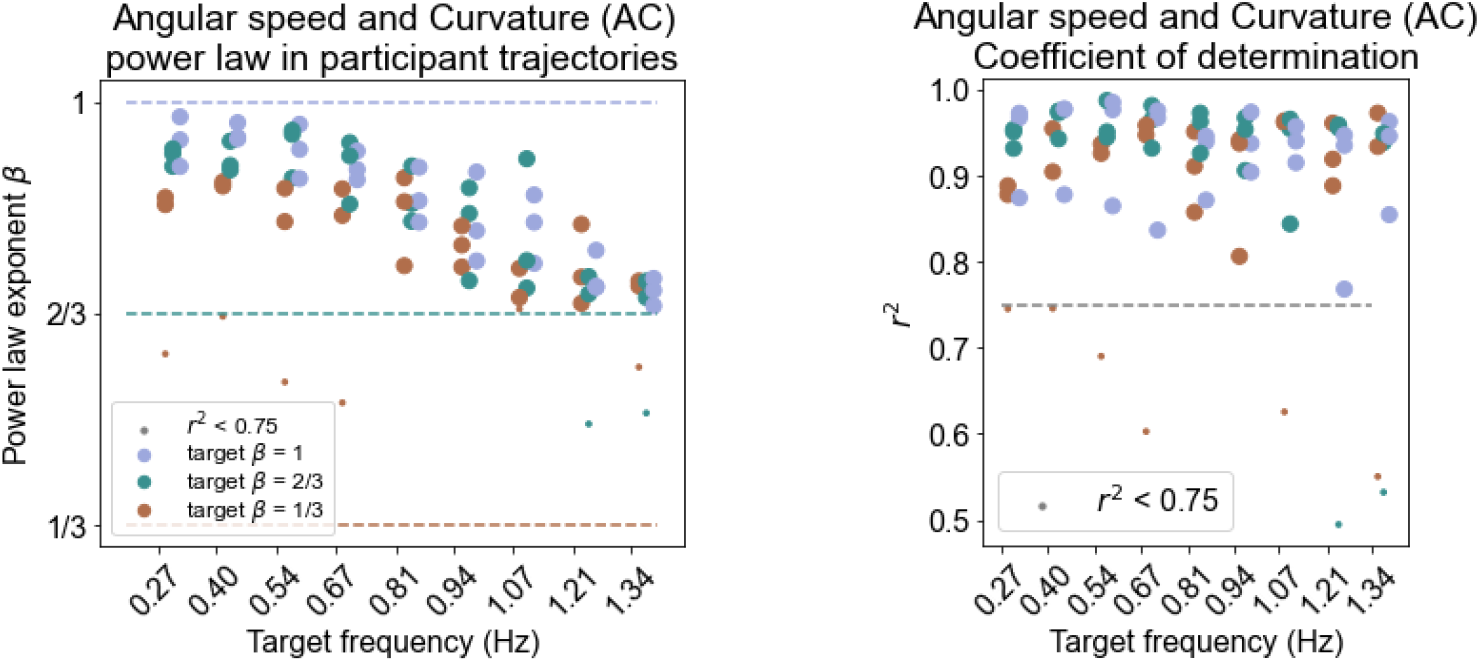
Using angular speed and curvature to estimate the speed-curvature power law on the same data as Figure 2, the r^2^ is higher than 0.75 for most trials, making the AC power law apparently much stronger than the VC power law.

Clearly, the VC and AC power laws are not equivalent (compare Figure 2 to Figure 3); the AC power law tends to be much stronger.

Figure 4 shows a common type of the differences in the estimated strength of the speed-curvature power law: when using angular speed, the power law is very strong while, for the same trajectory, tangential speed is very weakly related to curvature, apart from a slight tendency to slow down in more curved parts of the path.

**Figure 4.**
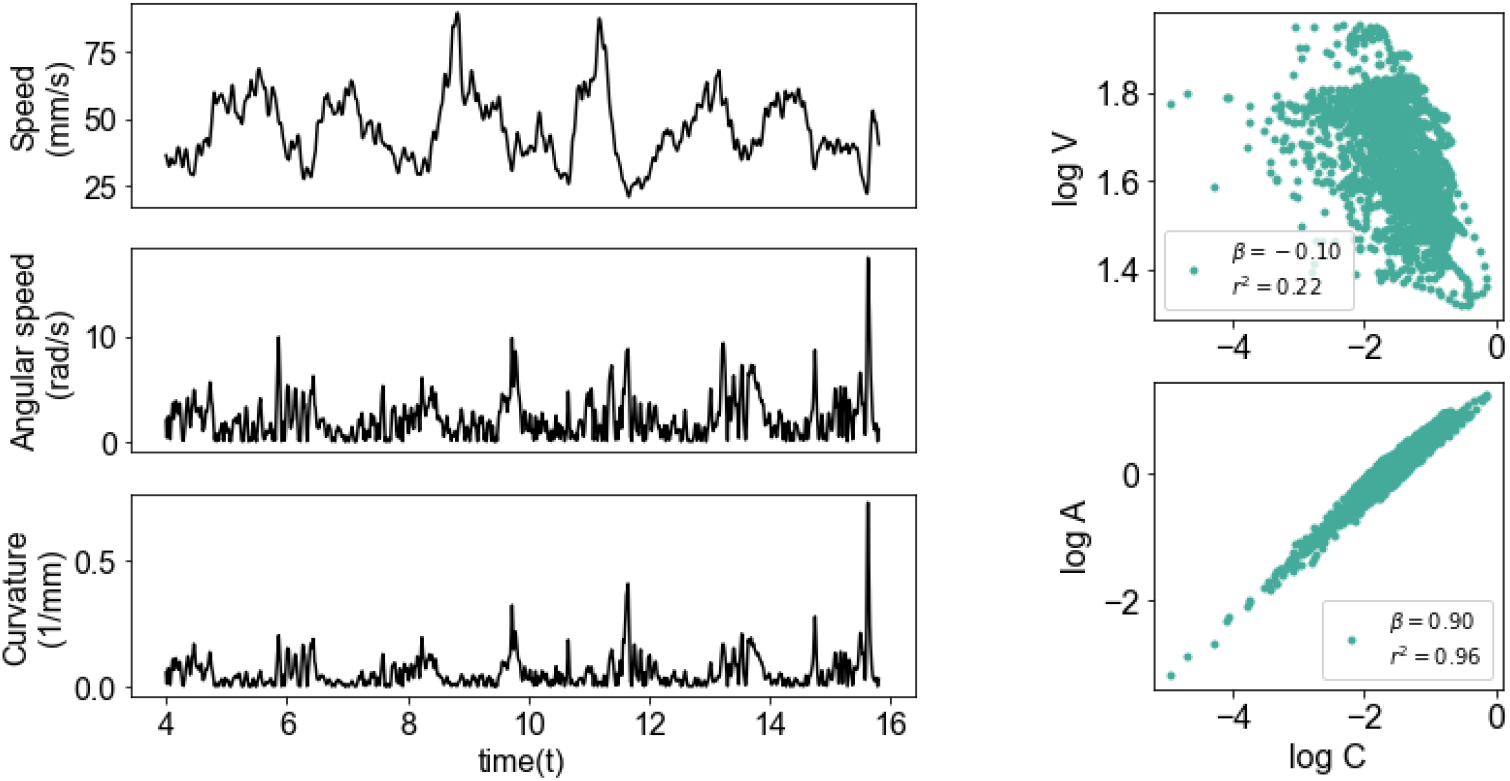
Example data from a single trial (target frequency = 0.27Hz), showing tangential speed, angular speed and curvatrue over time on the left. On the right, the VC power law is fairly weak (r^2^=0.22) showing that log tangential speed is hardly related to log curvature, while the AC power law is very strong (r^2^=0.96), since log angular speed is strongly correlated with log curvature.

In other words, there can be plenty of empirical cases where the speed V of the cursor is hardly related to the curvature, while the speed of ‘turning’, or the rate of change of direction, is strongly related to curvature. The following section explores why this is the case, using synthetic data.

## 3. SYNTHETIC DATA: Correlation between curvature and the two speeds Methods

In simple linear regression, the coefficient of determination r^2^ is equal to the square of Pearson’s correlation coefficient r. The Pearson’s correlation coefficient r is in turn defined as normalized covariance. To investigate the differences between the two power laws, here we focus only on the algebraic manipulations of correlations and covariances between the variables, and using simulations and random variables.

Random-speed elliptical trajectories were generated by first creating an elliptic trajectory with a given semi-minor and semi-major radius, frequency, and number of cycles, with a constant time-step. Next, a gaussian noise speed profile of the same duration as the trajectory is smoothed it with a low-pass butterwort filter at 10Hz cutoff. A time profile for the elliptic trajectory is created by dividing the distances between the points with the desired speeds to obtain from the random speed profile trajectory. Finally, using a cubic spline creates a constant time-sampling for the final elliptic trajectory (see the repository for the implementation: https://github.com/adam-matic/AngularSpeedPowerLaw)

### Results

According to equation (4), in the VC power law the variables correlated are log C and log V; according to equation (3), for the AC power law the variables correlated are log C and log A. However, we also know that A=CV, and therefore log A = log C + log V. In considering the AC power law, we are, in effect, correlating log C with log C + log V.

Letting Cov = covariance, a = log A, c = log C, and v = log V, we can write: Cov(c, a)

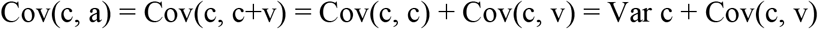

We can then see that Cov(c, a) will be dependent not only on the covariance of curvature and speed, but also on the variance of curvature. This means that the correlation coefficient will tend to be large when the curvature has a big variance relative to speed, regardless of the actual correlation between speed and curvature.

Consider two random normal variables x and y (Figure 5) and calculate the correlation coefficient between x and x + y, as an analogy to correlation of log C and log A. Letting Corr be equal to the Person’s correlation coefficient, obviously, the correlation r = Corr(x, x+y) is *not equivalent* to the Corr(x, y), which is zero in all four cases. Instead, if we keep the standard deviation of y stable, the Corr(x, x + y) increases with the standard deviation of x.

**Figure 5.**
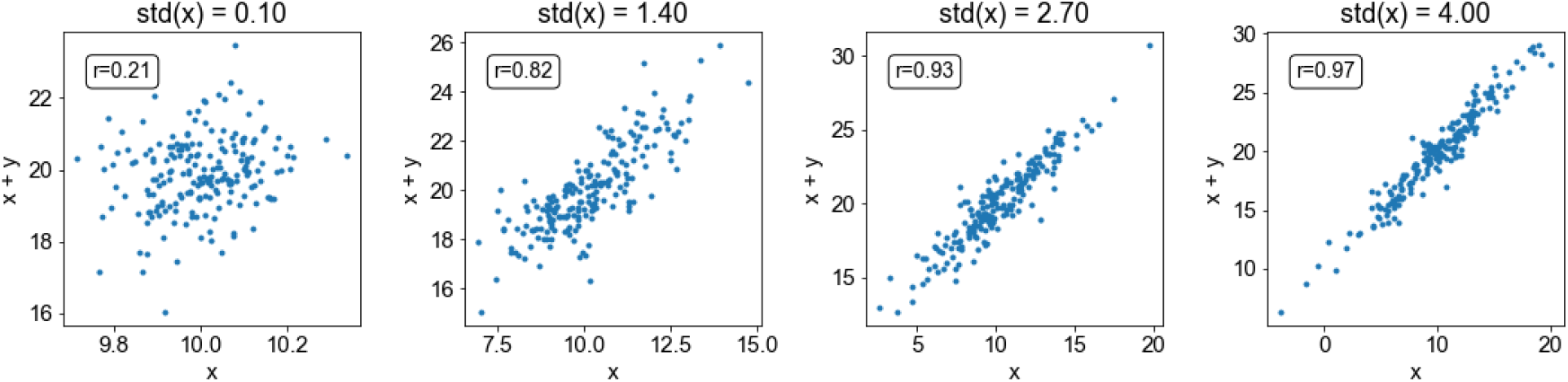
For two random normal variables x (mean 10, std in range [0.1, 4]) and y (mean 10, std 1), the variable x is not necessarily correlated to x+y, but it gets more strongly correlated as the variance of x increases.

The same effect of increasing the range of curvature can be illustrated with a random speed profile along a circular and along an elliptic trajectory. As shown on Figure 6A, the circular trajectory is designed to have a constant curvature profile and a random-smoothed speed profile. As expected, the correlation between logC and logV is very low, and so is the correlation between logC and logA. In the case of the ellipse (Figure 6B), the curvature profile has a bigger range, but the speed profile is identical to the circle. This single change was enough to increase the correlation between logC and logA to 1.

**Figure 6.**
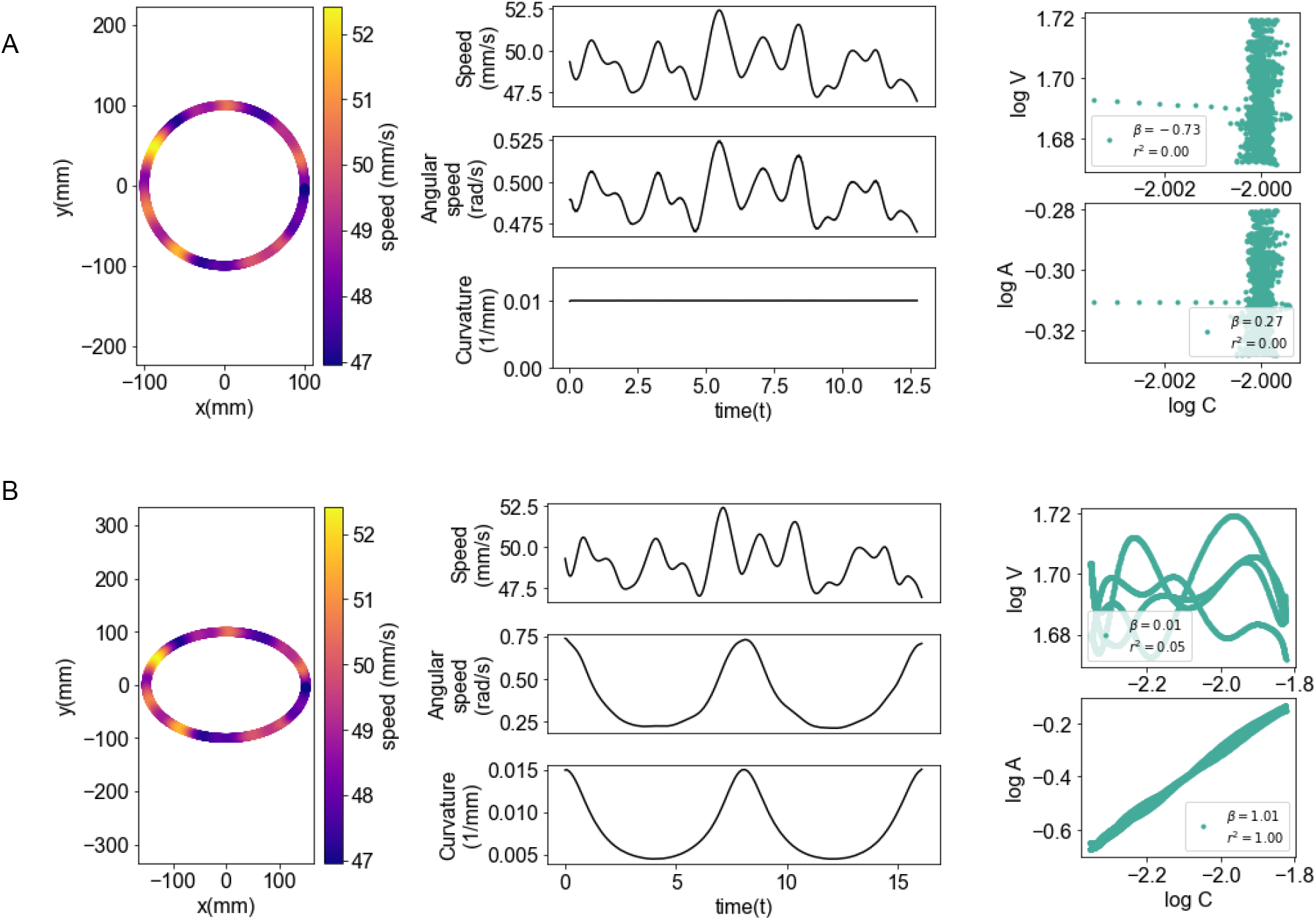
Correlation depends on the range of curvature in generated ellipses. **(A)** For a circular trajectory, radius of 100 mm, with a random-smoothed speed profile, both the logC-logV and logC-logA correlations are near zero. **(B)** For an elliptic trajectory, ra = 150 mm, rb =10 0mm, with the same speed profile, the logC – logV correlation is still low, while the logC-logA is very high.

We further explore this effect in Figure 7. We show how the Pearson’s correlation coefficient Corr(logC, logV) is near zero for all values the semi-major axis. This is expected, since the speed profile is randomized. However, for the correlation between angular speed and curvature, even a small increase in the semi-major axis, making the ellipse slightly eccentric, and making the curvature have slightly larger variance, increases the correlation Corr(logC, logA) to very high levels.

**Figure 7.**
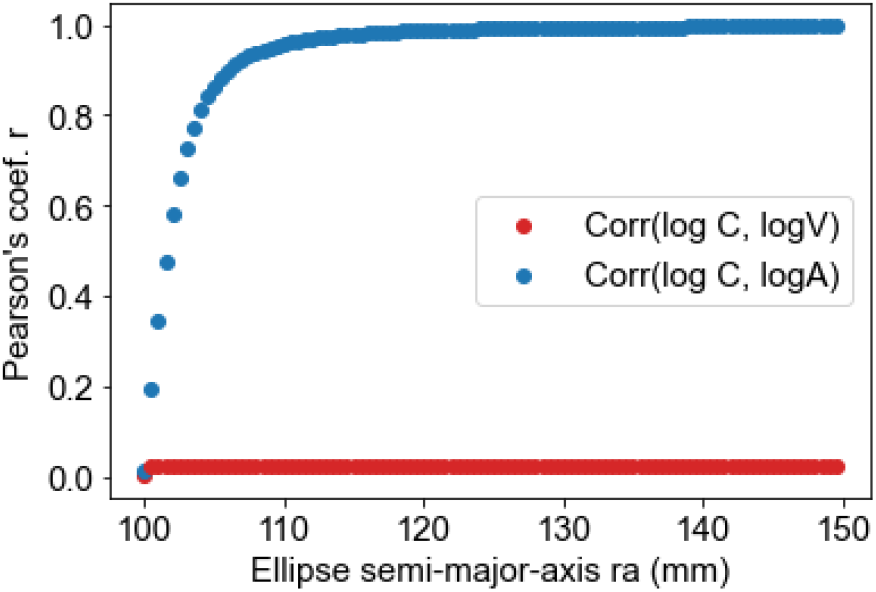
The correlation between logC and logA depends on the range of curvature, even in the absence of a speed-curvature power law. All ellipses had the same random-smoothed speed profile (see Figure 5), and the same semi-minor axis rb=100mm, while the curvature range was varied by increasing the semi-major axis. Note that, for instance, at 105 mm, which is 5% increase, the correlation is 0.87, and at 110 mm, which is a 10% increase, the correlation is 0.96.

**Figure 8.**
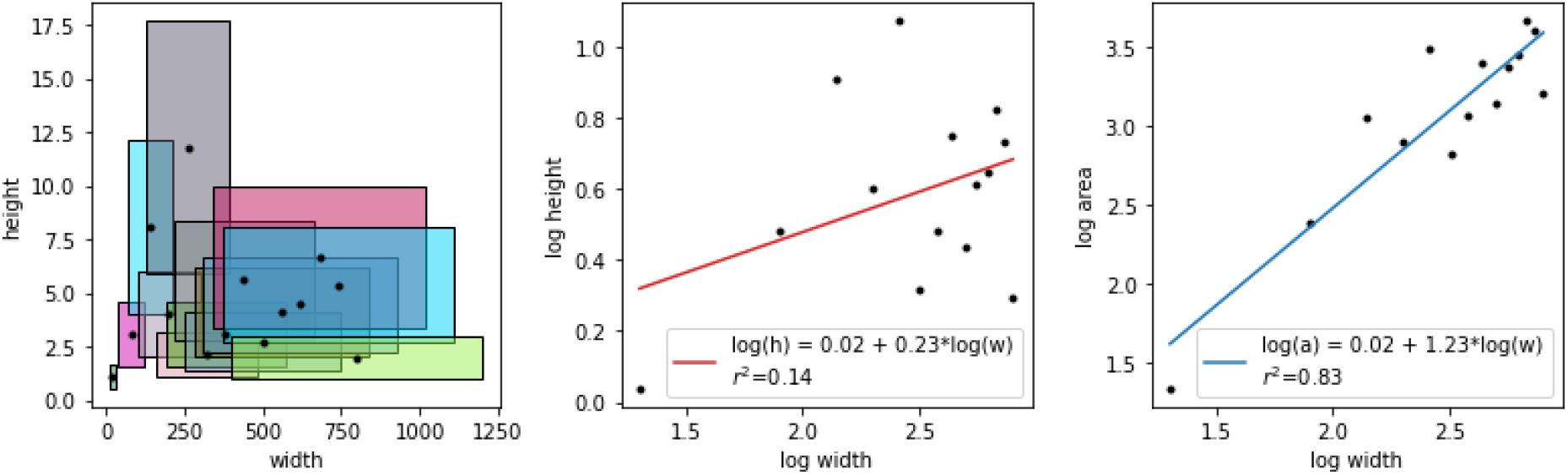
Figure 4. A set of rectangles of different sizes, random widths and heights. (Left) Widths and heights are uncorrelated. (Middle) Their logs are uncorrelated. (Right) Still, log(w) is strongly correlated to the log of area, log(a) because small-width rectangles tend to have small areas, and wide rectangles tend to have large areas; just like small curvature will tend to be associated to smaller angular speed, and larger curvature with larger angular speed. The exponent of the a-w power law is equal to the 1 + exponent of the w-h power law, repeating the AC and VC exponent relationship.

## 4. DISCUSSION

In this paper, I’ve analyzed the often-repeated statement in motor control research that tangential speed and angular speed are equivalent in estimating the speed-curvature power law from data. The analysis included empirical and synthetic data and the examination of the underlying algebra and mathematical relationships between the variables involved.

The analysis of empirical data from an experiment with human participants (Figures 2, 3 and 4) shows a huge difference between the two forms of the speed-curvature power law.

We found that fast drawing of ellipses, at about 1s of cycle time or faster, shows a strong speed-curvature (VC) power law, always with an exponent tending to βvc ≈-1/3 (Figure 2). Drawing equal-size ellipses at a lower frequency did not result in a strong VC power law. This correlation is measuring a genuine phenomenon in movement, an invariance that needs to be explained by theories of human motor control.

On the other hand, the correlation between angular speed and curvature was much stronger across the whole range of frequencies of drawing in the experiment (Figure 3). We can interpret this as a trivial correlation, since it reflects the tendency that small-curvature segments have smaller angular speed; a fact we know a priori from A = CV. The AC power law does not have an empirical origin; rather it is an artifact of the methods used to estimate it.

Conceptually, we could think of the correlation between angular speed and curvature as the correlation between widths and areas of rectangles in a set of rectangles of different sizes (area = width * height, or log area = log width + log height; in analogy to log V = log C * log A). Notably, analyzing the correlations between widths and heights of rectangles ignores the time dimension, just like the analysis of the speed-curvature power law in empirical data. The correlation is not necessarily there, but in most cases, if we have a sufficient range and variance of widths, the small-width rectangles can be associated with small areas, and large-width rectangles with large areas, regardless of their heights. The larger the range of widths, the stronger the correlation.

Although it is possible to measure empirical trajectories (Figure 4) or construct numerical trajectories (Figure 6) where angular speed and curvature are uncorrelated, in most empirically measured movement trajectories a great deal of the correlation between angular speed and curvature arises from the fact that the rate of change of direction increases in direct proportion to curvature. As illustrated, a high correlation between logA and logC in an elliptical trajectory can arise it complete absence of kinematic-geometric constraints (Figure 6B and Figure 7).

Ever since the pioneering works of Binet and Courtier (1883), Jack (1895), and a century latter Lacquaniti et al (1983), one goal of movement science is to understand why there is a relationship between movement speed and curvature. The main message of this manuscript is that when estimating the correlation between kinematics and geometry of movement, we should use mutually independent kinematic and geometric variables, such as tangential speed and curvature, and avoid angular speed, as it is often trivially correlated to curvature, and can lead to misinterpretations of the strength or the existence of the speed-curvature power law in empirical movement.

## Acknowledgements

I would like to thank Roberto Montanari, Javier Alegre Cortés and Alex Gomez-Marin for the valuable discussions and comments on earlier versions of the manuscript.

